# Invariant point message passing for protein side chain packing

**DOI:** 10.1101/2023.08.03.551328

**Authors:** Nicholas Z. Randolph, Brian Kuhlman

## Abstract

Protein side chain packing (PSCP) is a fundamental problem in the field of protein engineering, as high-confidence and low-energy conformations of amino acid side chains are crucial for understanding (and designing) protein folding, protein-protein interactions, and protein-ligand interactions. Traditional PSCP methods (such as the Rosetta Packer) often rely on a library of discrete side chain conformations, or rotamers, and a forcefield to guide the structure to low-energy conformations. Recently, deep learning (DL) based methods (such as DLPacker, AttnPacker, and DiffPack) have demonstrated state-of-the-art predictions and speed in the PSCP task. Building off the success of geometric graph neural networks for protein modeling, we present the Protein Invariant Point Packer (PIPPack) which effectively processes local structural and sequence information to produce realistic, idealized side chain coordinates using χ-angle distribution predictions and geometry-aware invariant point message passing (IPMP). On a test set of ∼1,400 high-quality protein chains, PIPPack is highly competitive with other state-of-the-art PSCP methods in rotamer recovery and per-residue RMSD but is significantly faster.

## Introduction

The myriad of complex functions facilitated by proteins as well as many intrinsic properties of proteins, such as folding and stability, are dependent on the interactions and conformations adopted by the protein’s amino acid side chains. Accurate modeling of side chains is therefore important for understanding structure-function relationships as well as designing new protein functions. The protein side chain packing (PSCP) problem has been traditionally formulated as a guided search over a library of discrete side chain conformations, or rotamers, given a protein backbone and its amino acid sequence^1^. There have been decades of research into and development of rotamer libraries that effectively capture the distribution of conformations observed for each amino acid in naturally occurring proteins^2–11^. Evaluating the favorability of individual rotamers in a residue’s environment often entails an energy function that models various physical phenomena such as hydrogen bonding and van der Waals interactions^12–17^. The final component of many traditional PSCP methods is a search strategy by which rotamers are sampled and evaluated across the entire protein^11,18–20^. Research in each of these individual components has led to the development of many traditional physics-based PSCP methods that have been successfully employed in a variety of applications.

Recently, the protein modeling field has been experiencing remarkable breakthroughs largely due to deep learning (DL) methods taking advantage of the growing amount of experimental protein data. Of particular significance, protein structure prediction networks, such as AlphaFold2 (AF2)^21^ and RoseTTAFold2 (RF2)^22^, have made large strides in predicting overall protein fold to near-experimental accuracy in many cases. While these methods produce coordinates for all heavy atoms in the protein and are, therefore, capable of side chain packing, there is no way to pre-specify and hold the backbone conformation fixed to mimic the PSCP task. On the other hand, DL-based methods designed specifically for PSCP have shown significant accuracy improvements over traditional approaches, while often being less time-consuming^23–29^. These methods either directly predict the location of side chain atoms^25,27^ or torsion angles from which the side chain can be reconstructed with idealized geometry^23,24,28^. Some of these methods rely on a rotamer library to select a specific conformation^25,26^ and some require subsequent post-processing to correct chemical and geometric violations and/or atomic clashes^27,29^.

Building off the recent success of graph neural networks for encoding and propagating structural information within proteins^27,30–33^, we present the Protein Invariant Point Packer (PIPPack) which can rapidly and accurately predict side chain conformations. After investigating the balance between data quality and dataset size, we trained our final model on non-redundant subset of the Protein Data Bank (PDB)^30^ to jointly predict binned χ dihedral angles for each residue, refining its previous predictions with recycling and explicitly incorporating, throughout the network, the geometry of the protein backbone through a novel message passing scheme. This message passing scheme can be viewed as a generalization of the invariant point attention (IPA) module introduced in AF2^21^ and, as such, is named invariant point message passing (IPMP). We demonstrate that incorporating IPMP for rotamer prediction provides a boost in performance over standard message passing and neighborhood-based IPA. Further performance improvements were obtained by fine-tuning the model with auxiliary losses and leveraging the joint knowledge of an ensemble of models. To improve the chemical and physical validity of the predictions, we further develop a simple resampling protocol that rapidly resolves most generated clashes. The training and inference code of our PyTorch^31^ implementation of PIPPack is publicly available on GitHub at https://github.com/Kuhlman-Lab/PIPPack.

## Methods

### Top2018 Dataset Preparation

The data used for training and evaluation was the Top2018 main chain-filtered dataset (v2.01, https://zenodo.org/record/5777651) created as described in Williams *et al*.^32^. Briefly, protein chains released prior to the start of 2019 that were solved by x-ray crystallography with a resolution of 2.0 or better were selected from the PDB^30^. They were subsequently filtered at a chain level to a set of chains with low MolProbity^33^ scores and few structural geometry outliers. Filtered chains were then clustered using MMseqs2^34^ to various sequence identity levels to reduce redundancy. For our models, we trained and evaluated using protein chains clustered at 40% identity. Next, the chains were subjected to filters applied to the main chain atoms (N, Cα, C, O, and Cβ) of each residue. Specifically, residues were removed from each chain if any atom under consideration had a high B-factor, geometry outlier, alternate location, steric clash, or did not agree with the experimental data. Finally, chains with more than 40% of its residues removed were discarded. In the end at 40% sequence identity, there were 10,449 clusters, of which 8,361 were used for training, 1,044 for validation, and 1,044 for testing. Finally, we removed any chains that had 40% sequence identity or more to a chain in the CASP13/14 test sets (see below, *n* = 439), as determined by MMseqs2’s easy-search workflow.

Note Williams *et al.*^32^ additionally created a full residue-filtered dataset wherein the same residue-level filters were applied to all heavy atoms in a residue. We decided to train on the main chain-filtered data primarily for two reasons. First, stricter filters in the full residue-filtered data results in fewer residues and, therefore, fewer training examples. Second, removal of residues whose side chain atoms do not pass the filters, but whose backbone atoms do results in loss in valuable training signals that may influence rotamer placement as, for instance, the N and O backbone atoms can participate in hydrogen bonding.

### BC40 Dataset Preparation

To further assess the balance between high-quality data and dataset size for PSCP, we additionally experimented with training models on the BC40 data^35^, which has been used recently for training data in PSCP methods^24,27^. This dataset consists of 36,970 protein chains released before August 2020 that are nonredundant at 40% sequence identity but have no other filters. It was originally constructed for the protein secondary structure prediction task^35^. We obtained the PDB code and chain identifier for each chain in the dataset, downloaded the coordinate files directly from the PDB, and extracted the appropriate chain. Prior to randomly splitting the data into training (90%) and validation (10%) sets, we removed any chains that had 40% sequence identity or more to a chain in the Top2018 test set (see above, *n* = 5,066) and the CASP13/14 test sets (see below, *n* = 736), as determined by MMseqs2’s easy-search workflow.

### CASP13 and CASP14 Test Set Preparation

In addition to the high-quality Top2018 test set, we evaluated our method on protein targets from the CASP13^36^ (*n* = 82) and CASP14^37^ (*n* = 64) competitions, like other recent PSCP methods^23,24,27,29^. Similar to the BC40 dataset, there are no structure-, chain-, nor residue-level filters, but this data was originally used to evaluate protein structure prediction methods. We acquired the PDB files for each target directly from the data archives of the CASP prediction center.

### Other PSCP Methods

To benchmark the performance of our method, we compared the results with four different previously released PSCP methods: Rosetta Packer^13,38^, DLPacker^26^, AttnPacker^27^, and DiffPack^24^. Rosetta Packer is the only non-DL based method considered here and is completely CPU bound. We interface with the packing protocol through PyRosetta^39^ (version 2021.36+release.57ac713), using the PackRotamersMover and the extra flags “-ex1 -ex2 -ex3 - ex4 -multi_cool_annealer 5 -no_his_his_pairE -linmem_ig 1”. DLPacker, AttnPacker, and DiffPack are all DL-based PSCP methods that take advantage of different neural network architectures and representations. The source code from the public release of these models (https://github.com/nekitmm/DLPacker, https://github.com/MattMcPartlon/AttnPacker, https://github.com/DeepGraphLearning/DiffPack) was downloaded along with the pre-trained model weights. Inference was performed in the standard protocol for each method, with the following notes: we used the “natoms” prediction order in DLPacker, we considered both AttnPacker with and without its post-processing step, and we also considered DiffPack with and without its confidence model. It should be noted that there is likely homology overlap between the Top2018 test dataset used for evaluation and the datasets used for training these models, so performance for these methods may be inflated.

### Architectural Considerations

Graph neural networks (GNNs) have shown remarkable promise in modeling proteins and have been successfully applied to various protein tasks, including fold classification^40,41^, property prediction^40–42^, fixed backbone sequence design^27,43–46^, and PSCP^24,27–29^. With the rationale that the specific side chain conformations are primarily dependent upon the local environment of the amino acid, we decided to model the PSCP problem with a GNN, wherein each residue is modeled as a node and is connected to its *k* nearest neighbors. Due to the various symmetries of proteins in 3D Euclidean space, special considerations must be taken in the formulation of the network, either preserving equivariance or invariance to global rotations and translations. Most networks for proteins that preserve equivariance due so by operating on equivariant features and predictions (e.g., coordinates) with specialized, equivariant neural network layers (e.g., SE(3) transformer^47^) to ensure that the effects of global transformations are propagated throughout the network^22,45,48^. Invariance, on the other hand, ensures that global transformations do not affect the network output and is often maintained using invariant features and predictions (e.g., relative orientations) and invariant layers (e.g., IPA)^21,46,49^. PIPPack is an invariant GNN that maintains its invariance through the choice of features, predictions, and layers (Fig. 1).

**Figure 1:**
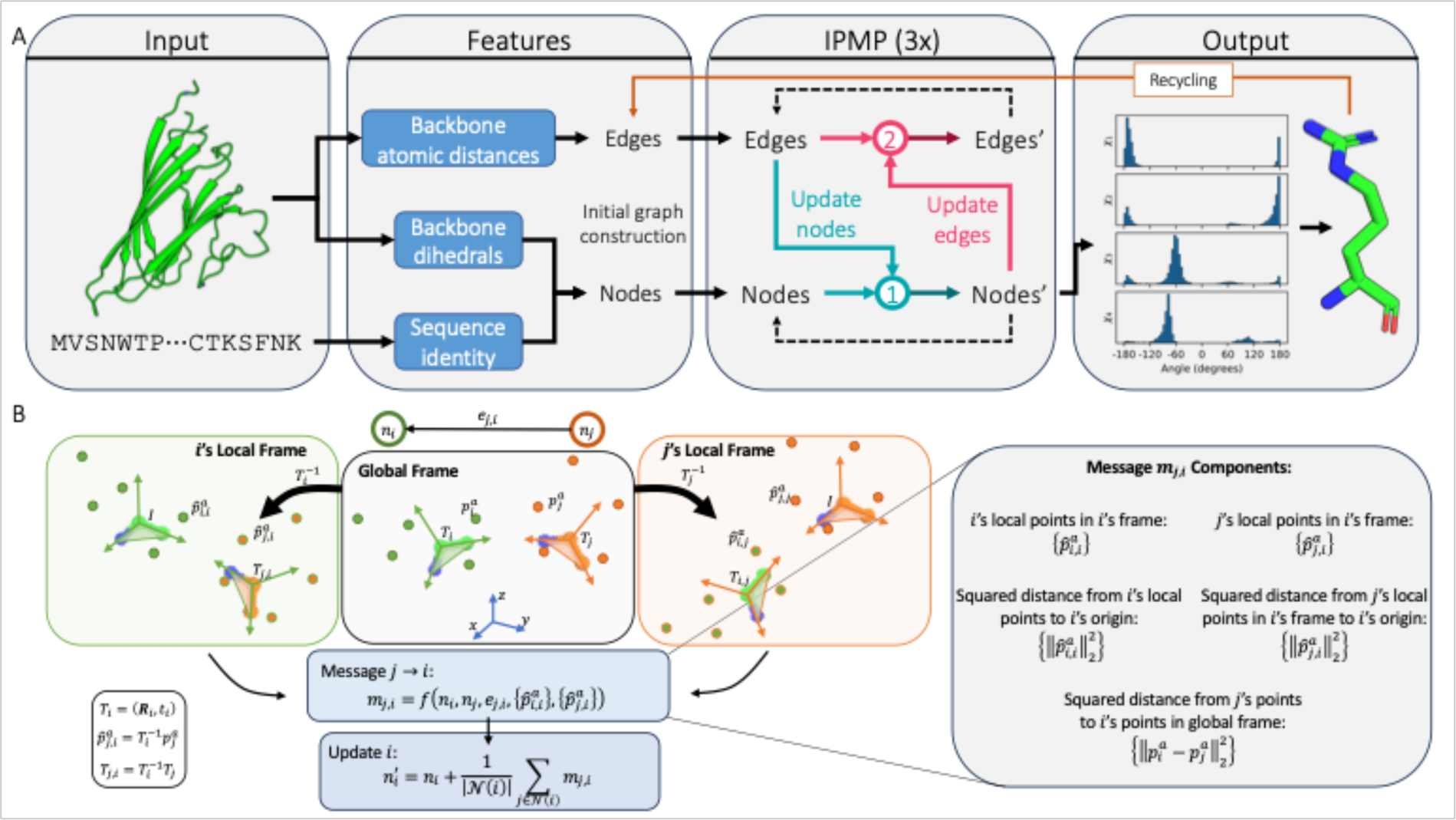
Architecture of PIPPack and invariant point message passing. (A) PIPPack is a graph neural network that processes protein backbone features (atomic distances, backbone dihedrals, and sequence) through geometry-aware message passing to iteratively refine and predict χ dihedral distributions for each residue. (B) PIPPack uses invariant point message passing (IPMP) to inject geometric information into each node and edge processing step. IPMP relies on invariant points that are produced in the local coordinate frame of each residue and transformed to obtain invariant features that depend on the protein backbone geometry.

### Initializing the protein graph

IPPack represents an input protein as a graph with a node for each residue and edges connecting nodes to their nearest neighbors (Fig. 1A). Node features include a one-hot embedding of the amino acid sequence and the sine and cosine of the backbone dihedral angles *ϕ, ψ*, and *ω*. The edges between residues contains a one-hot embedding of the relative sequence position and backbone-backbone atomic distances encoded with Gaussian radial basis functions (RBFs). Edges are formed between each residue and its *k* nearest neighbors (we use *k* = 30), determined by Cα-Cα distances. Note that all these features are invariant to global transformations. Additionally, for IPMP (see below, Fig. 1B), we obtain the rigid transformations that define the backbones of each residue, which are notably *equivariant* with respect to rigid transformations.

### Forming the predictions

The output of PIPPack is a packed protein structure, complete with all heavy atoms for each residue. We parameterize the PSCP task by predicting the χ dihedral angles for each residue, reconstructing the side chain with ideal bond lengths and angles. Most neural networks that predict side chain conformations via dihedrals perform a regression task on sin(χ) and cos(χ) for all χ angles, but we found improvements when we framed PSCP as a classification task by predicting the distribution across bins from [−*π*, *π*) (we used a bin width of 5°) as well as an offset value to precisely place the angle within the bin. Because the side chains are modeled with torsion angles, the prediction is also invariant. We note that this reframing also enables sampling from the predicted distributions to obtain some conformational diversity.

### Conditioning on previous predictions

AF2 utilized the concept of “recycling” whereby previous predictions were provided to the model, enabling multiple passes or attempts through the network^21^. This effectively conditions the model on its previous predictions, allowing them to be refined a few times. We incorporated recycling into PIPPack by augmenting the initial nodes and edges with features derived from the previously predicted side chains: node representations are updated with a sine and cosine encoding of the predicted χ angles and edge representations with additional side chain-backbone and side chain-side chain RBF-encoded distances. These previously predicted side chains are obtained by using the angle corresponding to the mode of the predicted distributions.

### Finetuning through discrete sampling

Inspired by the success of previous successful finetuning efforts^21,22^, we hypothesized that having loss terms that act on atomic coordinates to discourage producing clashes and unclosed proline rings would further bolster performance. Unfortunately, our reframing of PSCP as classification imposes serious difficulties in accurately backpropagating gradient signal from the coordinate losses through discrete samples from our predicted distributions. As concurrently introduced in Jang *et al.*^50^ and Maddison *et al.* ^51^, the Gumbel-Softmax (GS) trick can produce a differentiable sample from the predicted multinomial or categorical distribution by adding independently and identically distributed (iid) Gumbel noise to the model’s output logits^52^. Specifically, the GS distribution is defined as,

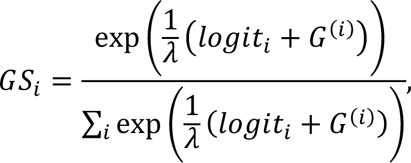

where *logit*_*i*_ is the log probability of the *i*th class, *G*^(i)^ is the *i*th iid sample from a standard Gumbel distribution, *λ* is a temperature parameter that controls the entropy of the distribution, and *GS*_*i*_ is the GS sample corresponding to the *i*th class. A hard or discrete sample from this distribution can be efficiently generated while preserving the gradient through the model logits. In a finetuning stage following standard model training, we employ this reparameterization trick (with *λ* = 1) for PIPPack’s χ angle distribution prediction to compute and train on two additional loss terms: (1) a clash loss that penalizes samples that result in atomic overlaps (determined with van der Waals radii), and (2) an unclosed proline loss that penalizes unclosed proline rings (determined by the Cδ-N bond length).

### Ensembling predictions

Due to the lightweight and one-shot nature of PIPPack, inference can be performed rapidly (Table 4). We hypothesized that predictions may benefit from combining knowledge from an ensemble of trained models, so we ensemble three randomly seeded models by simply averaging the predicted logits from each model before performing the softmax operation to obtain the final predicted probability distributions.

### Subsequent post-processing

When sampling from the predicted χ distributions, PIPPack can occasionally produce steric overlaps between atoms (Fig. S2, usually when forming hydrogen bonds, but not exclusively). Just as AttnPacker uses a post-processing step to correct potential violations in bond geometries and atom clashes, we reasoned that some form of minimization may reduce the clashes in PIPPack’s predictions. We experimented with applying Rosetta’s MinMover protocol (referred to as PIPPack+RM) that reduces the energy of Rosetta’s energy function (ref2015) by gradient descent and manipulating specific degrees of freedom (in our case, side chain torsional angles) and the same post-processing procedure (referred to as PIPPack+PP) as AttnPacker which applied gradient-based minimization to reduce clashes while not straying too far from the original torsion predictions (Table 2). Because the side chains are constructed by sampling the predicted distributions, we also created a resampling protocol for PIPPack (referred to as PIPPack+RS) that analyzes the sampled structure to identify clashing residues which then have their χ angles resampled using Markov chain Monte Carlo (MCMC) with a Metropolis criterion. Because sampling from the predicted probability distributions with low temperature results in the best performance but low diversity, our resampling procedure gradually raises the temperature to try to balance sampling high-probability conformations while introducing conformational diversity. In this resampling protocol, we additionally resample χ angles for prolines residues if the proline is not closed.

### Exchanging information with message passing

Graph neural networks often extract and process information from example graphs by performing convolution or passing messages between neighboring nodes to update node and/or edge representations^53^. This complete update step is usually made up of three functions, that is, to perform a node update for *n*_*i*_, we perform

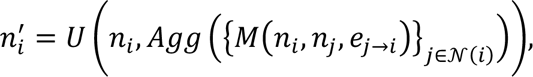

where *M* is the message function that computes the message between nodes *i* and *j* based on node information *n*_*i*_, *n*_*j*_ and directed edge information *e*_*j→i*_, *Agg* is a permutation-invariant aggregation function (e.g., mean or sum) that combines messages from all neighbors *j* in the neighborhood *N*(*i*) of node *i*, and *U* is an update function that computes the new node state based on the previous state and the aggregated messages. The neighborhood *N*(*i*) is defined as all the nodes connected to node *i*.

We explore the use of three types of message passing layers: a standard messaging passing layer (MPNN), a neighborhood-based IPA layer (IPA), and a novel invariant point message passing layer (IPMP, see next section). The update function of the MPNN layer used in this paper is given

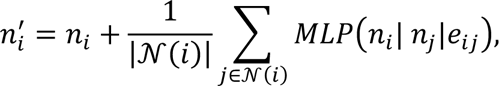

where |*N*(*i*)| is the number of neighbors in *N*(*i*), *MLP* is a multi-layer perceptron, and ⋅ | ⋅ denotes the concatenation operation. The update function of the IPA layer used in this paper is given as

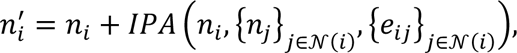

where *IPA* is the invariant point attention module introduced in AF2 (Algorithm 22 in the supplementary information of AF2^21^) but restricted to computing attention on the |*N*(*i*)| neighbors of node *i* rather than the entire set of nodes.

### Invariant Point Message Passing (IPMP)

The protein structure prediction network AF2 introduced a geometry-aware node representation update termed “invariant point attention” (IPA)^21^. This operation performs attention across each node (residue) biased by information contained in the edges and geometric proximity. To capture this geometry awareness, IPA utilizes rigid transformations *T*_*i*_ = (***R**_*i*_, t* →_*i*_), where ***R***_*i*_ is a rotation matrix and *t* →_*i*_ is a translation vector, that represent the backbone of each residue *i* and places “invariant points” (points in the local frame of *i*). This information is aggregated via an edge- and geometry-biased attention mechanism and used as an update for each node. Note that the rigid transformations are not invariant with respect to global transformations and, therefore, must be applied appropriately to maintain invariance. In AF2, the protein is essentially represented as a densely connected graph and, therefore, messages come from every pair of nodes, but IPA can easily be adapted to form messages within local neighborhoods. In the message passing framework, IPA uses a message function that consists of geometry- and edge-biased attention.

While attention is a powerful and performant operation, it may be useful to be able to consider other functions that operate in a geometry-aware manner like IPA but without the attention. To this end, we generalize IPA to “invariant point message passing” (IPMP) wherein the message function becomes some invariant function of the connected nodes, the edge between them, and their rigid transformations, i.e., *M*(*n*_*i*_, *T*_*i*_, *n*_*j*_, *T*_*j*_, *e*_*j→i*_) (Fig. 1B). We specifically experiment with a message function that concatenates the node embeddings *n*_*i*_and *n*_*j*_, the edge embedding *e*_*j→i*_, and five components derived from local points for each node and their rigid transforms. That is, first, each node computes *N_points_* invariant points from its node representation with a learnable function 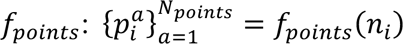 (for this, we simply use a linear layer). The five additional components are computed by combining the invariant points and the rigid transforms of each residue:

- Invariant points in node *i*’s local frame: 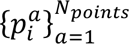
- Squared distance between node *i*’s origin and node *i*’s invariant points: 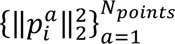
- Invariant points from node *j* in node *i*’s local frame: 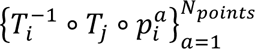
- Squared distance between node *i*’s origin and node *j*’s invariant points in node *i*’s local frame: 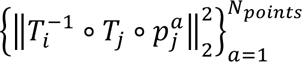
- Squared distances between points in global frame: 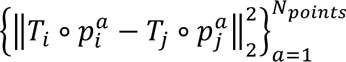

These components are concatenated with the node and edge embeddings and are processed with an MLP to obtain the message *m_ij_*. The same aggregation and update functions as shown for the MPNN layer are then applied to create the new node representation *n’_i_*. Note that each time the transformations *T*_*i*_ are applied, it is in an invariant manner (e.g., distance/norm calculation). We also note that the specific form of the message function *M* need not be constrained to all or any of the five components listed above, so long as the message function remains invariant to global transformations. We leave the further customization of this function (which may be task-dependent) to future work.

### Training PIPPack

PIPPack was trained using PyTorch^31^ until convergence with early stopping on validation χ perplexity and the Adam optimizer with learning rate schedule described in Vaswani *et al* ^54^. Using an NVIDIA A100 80G GPU, the network trained for approximately 5.5 days (BC40 dataset) or 14 hours (Top2018 dataset) using a random contiguous crop size of 512 residues and a batch size of 32 chains. The final model is relatively lightweight with about 1.9 M learnable parameters but can be run as an ensemble of 3 randomly seeded models. As mentioned in above, we finetuned PIPPack using an additionally clash loss and unclosed proline loss that act on a GS sample from the predicted χ distributions. During finetuning, we train to convergence using the Adam optimizer with a fixed learning rate of 1e-8 (another 5 days on BC40 data).

### Evaluation of Performance

Following other PSCP methods, we evaluate the performance of our method (and other methods) on our Top2018 test set using residue-level root mean squared deviation (RMSD), χ dihedral angle mean absolute error (MAE), and rotamer recovery (RR). Residue-level RMSD is determined by aligning the backbone atoms (N, Cα, C, and O) of the predicted and ground-truth residues and computing the RMSD over the side chain heavy atoms (including Cβ), and we report the mean RMSD value (in Å) over all residues in the test set. χ MAE (in °) is computed by determining the absolute error for each χ_*i*_ and averaging over all χ_*i*_ in the dataset. RR is the percentage of recovered rotamers within the dataset, where a rotamer is considered recovered if *all* the predicted χ_*i*_ for a particular residue are within 20° of the native χ_*i*_. These metrics are further stratified across amino acid type and the different centrality levels: all, core, and surface. A residue is considered in the core if the number of neighboring residues (determined by Cβ-Cβ distance < 10 Å) is at least 20, whereas it is considered on the surface if there are at most 15 neighbors. Additionally, we report the mean clashscore and rotamer evaluations, both determined via MolProbity^33^. Clashscore refers to the number of serious steric clashes (atoms with overlap of van der Waals radii > 0.4 Å) per 1000 atoms, whereas rotamer evaluations determine if a specific rotamer is considered “favored”, “allowed”, or an “outlier” based on the statistics of occurrences within the PDB.

## Results

### PIPPack Ablation Studies

To investigate the contributions from different architectural decisions, we systematically removed components from PIPPack and retrained our network on the Top2018 dataset. Specifically, we explored the importance of the χ angle representation (discretized bins vs sine and cosine), the benefit of geometry-aware updates (i.e., MPNN layers vs IPA layers vs IPMP layers), the role of iterative prediction refinement via recycling, result of finetuning, and the effects of ensemble predictions from multiple models (Fig. 2).

**Figure 2:**
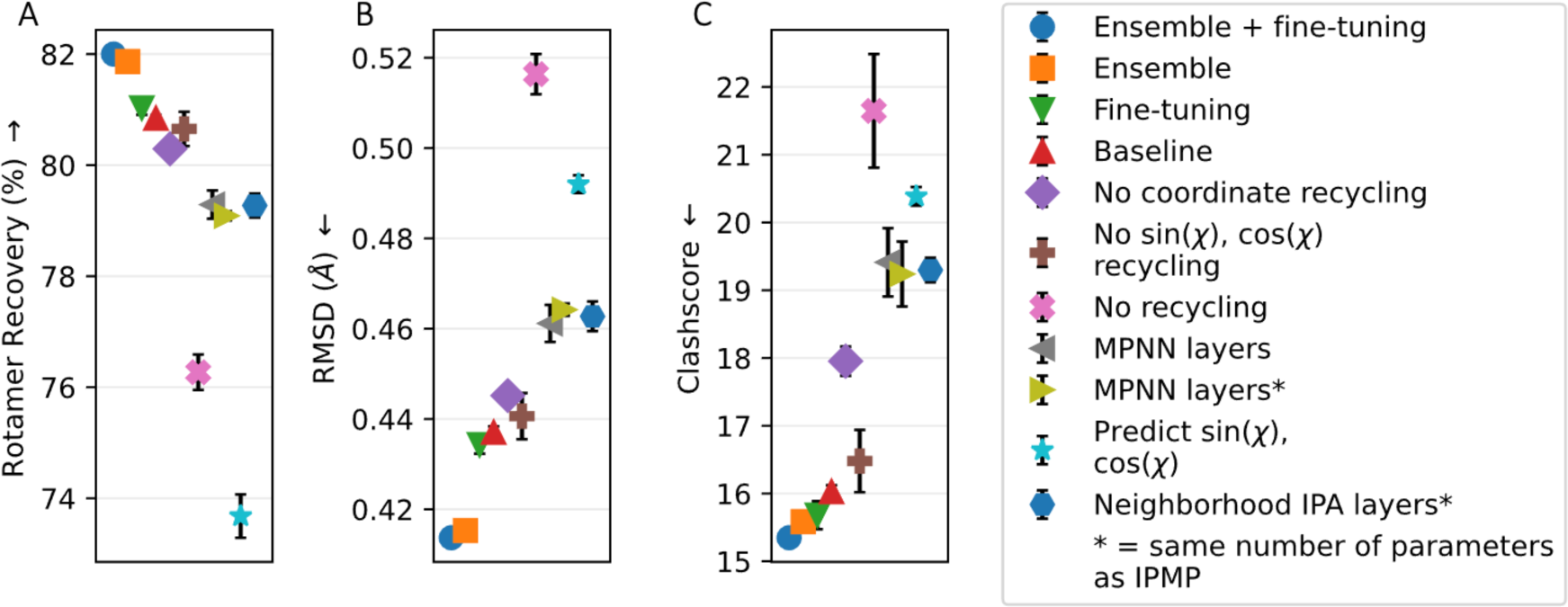
PIPPack ablation studies. Components were systematically removed from PIPPack to assess their contributions to its performance, specifically with respect to (A) rotamer recovery, (B) root mean squared deviation (RSMD), and (C) clashscore. The baseline model refers to PIPPack trained on Top2018 with binned χ prediction, 3 recycles, and IPMP layers.

One of the two largest contributors to PIPPack’s success was the transformation of the PSCP problem from regression to classification, affecting the model’s RR by more than 7%. Concretely, for each χ_*i*_, this transforms the prediction from the continuous sin(χ_*i*_), cos(χ_*i*_) target to predicting discrete probabilities for each bin across [−*π*, *π*). This resulted in additional benefits in terms of RMSD (∼0.055 Å) and clashscore (∼4.35). For these evaluations, the mode of the final predicted distributions was used as the output χ angle.

Recycling, the other major contributor, provided an improvement of about 4.5% in RR, 0.08 Å in RMSD, and 5.6 in clashscore by enabling PIPPack to iteratively refine its previous predictions. Interestingly, although recycling doesn’t improve RR as much as the classification reframing, it provides even larger benefits in terms of RMSD and clashscore. Between the two types of recycled information, the coordinates of the predicted side chains are more beneficial than the sine and cosine of the predicted χ angles. Providing both these features yields a model with slightly better performance. While the model was trained with a specific number of recycling iterations in mind, the actual sampling procedure can perform an arbitrary amount of recycles. The default protocol utilizes the number of recycles that the model was trained for (in our case, 3 recycles), but it has been shown in other models that incorporate recycling, such as AF2, that additional recycling iterations can have some benefits in terms of the prediction accuracy^55,56^. To evaluate this effect for PIPPack, we sampled rotamers for the Top2018 test set from the baseline model while varying the number of recycling iterations from 0 to 6 (Fig. S1). Increasing the number of recycling iterations appears to have little effect on the mean performance metrics past the default value, suggesting a limit to the model’s refinement capabilities. Increasing recycling iterations also has the downside of increasing runtimes, requiring *n* + 1 passes through the network for *n* recycles. Interestingly, however, PIPPack’s performance shows the greatest improvement as the number of recycles increases from 0 to 1, suggesting that model inference with just a single recycle may strike a balance between speed and prediction quality if necessary.

Changing the types of layers used within the model from IPA or MPNN layers to IPMP layers led to an additional modest performance boost (1.5-1.75% for RR, 0.024-0.027 Å for RMSD, and 3.2-3.4 for clashscore). Moreover, ensuring roughly the same parameter count between these variant models suggests that IPMP’s performance isn’t simply due to larger model capacity. To accommodate similar parameter counts between layers, we increased the number of channels inside the MPNN layer and reduced the number of heads within the IPA layer. The success of IPMP layers suggests that explicit incorporation of geometry-aware updates without attention-weighted messages provides better inductive reasoning over the protein structure.

To improve past the baseline model, we employed two techniques: finetuning and ensemble prediction. Further training of the model with the additional auxiliary losses improved PIPPack over the baseline by about 0.2% in RR, 0.003 Å in RMSD, and 0.3 in clashscore. Without compromising too much speed, ensembled PIPPack benefits by averaging the predicted χ distributions of several trained models. Ensembling is accomplished via averaging the outputted logits (prior to the softmax computation for the probabilities) from three randomly seeded versions of our model. The combined knowledge enabled improvements over the baseline across rotamer recovery, RMSD, and clashscore of about 1%, 0.022 Å, and 0.5, respectively. Furthermore, ensembling the predictions of the finetuned models results in improvements of 1.16% in RR, 0.023 Å in RMSD, and 0.68 in clashscore over the baseline.

### Determining the Importance of Data Quality and Dataset Size

To evaluate the effect of training on data subjected to different quality filters and in datasets of different size, we trained models (without finetuning) on the Top2018 data and the BC40 data. Moreover, we experimented with an additional quality filter, applied directly at runtime: B-factor filters, wherein any χ angle that depends on a side chain atom with B-factor > 40 Å^2^ is discarded. This filter and the two datasets result in four training-set regimens: Top2018 data with and without B-factor filter (Top2018-BF and Top2018) and BC40 data with and without B-factor filter (BC40-BF and BC40). These models trained with each regime were then evaluated (in triplicate) on the Top2018 test set and the CASP13/14 test sets. To assess another dimension of dataset quality, we report performance on our test sets using residues filtered such that the side chain atoms have low B-factors (< 40 Å^2^).

As seen in Table 1, applying B-factor filters to the test sets results in fewer total residues for consideration, with the largest differences occurring in the CASP datasets wherein most of the residues (> 60%) are filtered out. This filtered subset of the test sets represents the residues whose side chain conformations are reasonably reliable and, as such, likely comprises a better estimate of the true performance of PSCP methods. Removing the high-B-factor dihedrals, however, likely biases the distribution of residues towards core residues, which are generally more rigid due to well-defined interactions with their neighbors and are, intuitively and objectively, easier to predict. Interestingly but not unexpectedly, when we apply the B-factor filters, the metrics improve and have smaller deviation between test sets.

**Table 1:** Impact of B-factor filters on training and testing data.

| Training Set | Test Set |  |  |  |  |  |  |  |  |  |  |  |
| --- | --- | --- | --- | --- | --- | --- | --- | --- | --- | --- | --- | --- |
|  | Top2018 |  | Top2018-BF |  | CASP13 |  | CASP13-BF |  | CASP14 |  | CASP14-BF |  |
|  | RMSD<br>(Å) ↓ | RR<br>(%) ↑ | RMSD<br>(Å) ↓ | RR<br>(%) ↑ | RMSD<br>(Å) ↓ | RR<br>(%) ↑ | RMSD<br>(Å) ↓ | RR<br>(%) ↑ | RMSD<br>(Å) ↓ | RR<br>(%) ↑ | RMSD<br>(Å) ↓ | RR<br>(%) ↑ |
| Top2018 | 0.493 | 77.36 | 0.433 | 81.11 | 0.741 | 61.51 | 0.594 | 71.05 | 0.910 | 51.08 | 0.585 | 68.98 |
| Top2018-BF | 0.495 | 77.28 | 0.434 | 81.05 | 0.751 | 61.13 | 0.603 | 70.81 | 0.920 | 50.82 | 0.589 | 68.95 |
| BC40 | **0.472** | **78.77** | **0.412** | **82.53** | **0.660** | **65.96** | **0.535** | **74.77** | **0.816** | **55.43** | **0.539** | **72.06** |
| BC40-BF | 0.487 | 77.90 | 0.425 | 81.75 | 0.704 | 63.74 | 0.565 | 73.06 | 0.872 | 53.05 | 0.562 | 70.74 |
| **Number of Rotamers** | 259,911 |  | 239,167<br>(92.02%) |  | 18,020 |  | 7,199 (39.95%) |  | 13,356 |  | 4,449 (33.31%) |  |

Training models with B-factor cutoffs appears to only decrease the overall performance of the method, regardless of the training dataset used. This might be explained by the number of residues that can serve as training data in each dataset. When B-factors are applied to the BC40 dataset (*n* = 7,130,551 residues), 50.16% or 3,576,396 residues are removed. For the Top2018 dataset, 7.36% are removed (199,920 of 2,715,530 residues). The performance metrics correlate with the dataset size, suggesting that amount of trainable dataset is more important than ensuring that the data is high quality.

Across testing datasets, the models trained on the BC40 dataset performed about 1.25 – 4% better in RR than those trained on Top2018. The BC40 training set contains about 2.5 times more chains and more rotamers than Top2018 but with much less stringent quality filters, reinforcing the importance of dataset size. Differences between the BC40-and Top2018-trained models are most apparent in the CASP13/14 test sets, but due to the 30-50 times more rotamers, we consider the results from the Top2018 datasets to be more accurate and robust estimates of the true model performance. Based on this analysis and the ablation study, PIPPack trained and finetuned on BC40 data without B-factor filters is the model we pursue in subsequent analyses and comparisons to other methods, and we refer to this model as PIPPack for the rest of the paper (unless otherwise noted).

### Post-Processing of PIPPack Predictions

Although PIPPack can rapidly produce accurate side chains, it occasionally produces steric overlaps between atoms (usually when forming hydrogen bonds, but not exclusively, see Fig. S2) and unclosed proline residues. Just as AttnPacker uses a post-processing step to correct potential violations in bond geometries and atom clashes, we reasoned that some form of minimization may reduce these issues in PIPPack predictions. We experimented with applying Rosetta’s MinMover protocol (referred to as PIPPack+RM) that reduces the energy of Rosetta’s energy function (ref2015) by gradient descent and manipulating specific degrees of freedom (in our case, side chain torsional angles) and the same post-processing procedure (referred to as PIPPack+PP) as AttnPacker which applied gradient-based minimization to reduce clashes while not straying too far from the original torsion predictions (Table 2). Both post-processing procedures reduce PIPPack’s overall RR performance, but only the MinMover improves RMSD. AttnPacker’s post-processing, however, improves the clashscore more than Rosetta MinMover, presumably because the minimization objective function in Rosetta contains more terms than just a repulsive clash energy. These energy terms may also provide some explanation as to the slight improvements in MAE of longer side chains, specifically χ_3_ and χ_4_, and RMSD.

**Table 2:** Post-processing of PIPPack predictions on Top2018 test set.

| Method | RMSD (Å) ↓ | | | $\chi$ MAE (°) ↓ | | | | RR (%) ↑ | | | Clashscore ↓ | Post-processing time (avg. s/protein) |
| --- | --- | --- | --- | --- | --- | --- | --- | --- | --- | --- | --- | --- |
| | All | Core | Surface | $\chi_1$ | $\chi_2$ | $\chi_3$ | $\chi_4$ | All | Core | Surface | | |
| PIPPack | <u>0.412</u> | 0.310 | <u>0.506</u> | **9.69** | **14.93** | <u>30.66</u> | 41.63 | **82.53** | **88.89** | **75.94** | 14.50 | --- |
| PIPPack + RM | **0.404** | **0.298** | **0.500** | 10.11 | 15.47 | **30.45** | **41.20** | <u>81.99</u> | 88.31 | <u>75.49</u> | 12.68 | 19.07 |
| PIPPack + PP | 0.418 | 0.315 | 0.512 | <u>9.82</u> | 16.04 | 32.31 | <u>41.58</u> | 81.74 | 88.12 | 75.18 | <u>10.31</u> | 8.38 |
| PIPPack + RS | 0.413 | <u>0.309</u> | 0.509 | 9.86 | <u>15.26</u> | 31.17 | 42.13 | 81.93 | <u>88.39</u> | 75.29 | **9.95** | 0.45 |

In addition to applying these minimization protocols, we also experimented with a resampling algorithm that simply identifies clashing and unclosed proline residues and resamples the χ distributions for those residues with MCMC and a Metropolis criterion. In comparison to the previous two approaches, resampling leads to the largest improvement of clashscore with relatively minor effects on the other metrics. Another major benefit of this resampling protocol is that no additional model evaluations or gradient calculations are necessary, resulting in minimal increase in runtime (Table 2).

### Performance Comparison with PSCP Methods

We sought to evaluate PIPPack’s performance in the context of other successful PSCP methods, specifically Rosetta Packer^13,38^, DLPacker^26^, AttnPacker^27^, and DiffPack^24^. DLPacker^26^ is a PSCP method that sequentially captures the local environment of each residue by performing 3D convolutions, predicts a probability density for the location of side chain atoms for a residue, and then selects a rotamer that fits into the predicted density from a rotamer library. Because DLPacker operates on each residue within the protein one at a time, there can be different orders by which the rotamers are sampled. We follow the recommendation by Misiura *et al.*^26^ that assigns rotamers sequentially from the most crowded residues to the least crowded residues, as this order serves as a compromise between speed and quality.

AttnPacker^27^ is an attention-based GNN that processes the protein backbone through equivariant updates to finally predict the locations of side chain atoms all at once. Because the network is predicting coordinates of all side chain atoms simultaneously, AttnPacker sometimes violates chemical bond geometries and produces atomic clashes, therefore requiring a post-processing step to idealize the side chains and reduce these violations. Moreover, McPartlon *et al* ^27^ also introduced an inverse folding variant of AttnPacker that designs an amino acid sequence and packs the rotamers. As the two variants of AttnPacker perform similarly for the PSCP task, we only consider the packing variant with and without subsequent post-processing in our comparison.

DiffPack^24^ is a diffusion-based method that iteratively denoises the torsional distribution of χ angles, utilizing a series of SE(3)-invariant GNNs as score networks to autoregressively build up each side chain. Note that unlike any of the other methods benchmarked here, DiffPack is the only network that applies diffusion-based generative PSCP and builds the side chain of each residue one χ angle at a time. A confidence-aware version of DiffPack uses another network to predict error in the modelled side chains and allows for sampling of multiple trajectories and combining predictions that are the most confident. We benchmark both DiffPack with and without the additional confidence model. As mentioned in the previous section, steric clashes generated by PIPPack are reduced when post-prediction minimization is applied, so we additionally consider PIPPack with resampling (PIPPack+RS).

As above with the evaluation of the datasets, we evaluate the packing solutions on the Top2018 test data with low B-factors in residue RMSD and RR but also consider χ MAE, clashscore, and rotamer evaluations, stratifying some of these metrics by centrality level (Table 3). With respect to the χ angle performance metrics (χ MAE and RR), ensembled PIPPack and PIPPack+RS outperforms all the other PSCP methods. In terms of RMSD, PIPPack performs quite competitively, achieving top-2 RMSD with the ensembled version. PIPPack does however rely on the resampling procedure to obtain less clashes than most other methods.

**Table 3:** Protein side chain packing results on B-factor-filtered Top2018 test set.

| Method | RMSD (Å) ↓ | | | $\chi$ MAE (°) ↓ | | | | RR (%) ↑ | | | Rotamer Outliers<sup>†</sup> (%) | Clashscore<sup>†</sup> ↓ |
| --- | --- | --- | --- | --- | --- | --- | --- | --- | --- | --- | --- | --- |
| | All | Core | Surface | $\chi_1$ | $\chi_2$ | $\chi_3$ | $\chi_4$ | All | Core | Surface | | |
| Rosetta Packer | 0.589 | 0.423 | 0.733 | 18.28 | 22.35 | 37.22 | 45.41 | 71.97 | 80.85 | 63.24 | 0.07 | 12.40 |
| DL Packer | 0.443 | 0.340 | 0.535 | 12.17 | 19.62 | 39.34 | 56.85 | 75.19 | 81.71 | 68.77 | 1.17 | 10.04 |
| AttnPacker | 0.519 | 0.450 | 0.574 | 12.93 | 29.68 | 35.82 | 45.35 | 65.68 | 70.43 | 61.19 | 4.01 | 21.85 |
| AttnPacker+PP | 0.440 | 0.349 | 0.520 | 10.95 | 20.62 | 40.67 | 45.18 | 73.80 | 79.78 | 67.91 | 2.29 | 11.17 |
| DiffPack | 0.407 | 0.308 | 0.490 | 11.47 | 17.45 | 34.21 | 43.51 | 79.72 | 85.55 | 74.02 | 0.55 | 9.84 |
| DiffPack +Confidence | **0.347** | **0.256** | **0.426** | 9.59 | 15.44 | 30.72 | <u>39.43</u> | 82.52 | 88.13 | <u>76.98</u> | 0.43 | **6.74** |
| PIPPack<sup>‡</sup> | 0.406 | 0.305 | 0.498 | 9.52 | 14.72 | 30.13 | 40.36 | 82.92 | 89.25 | 76.37 | 0.17 | 14.25 |
| PIPPack+RS<sup>‡</sup> | 0.407 | 0.304 | 0.500 | 9.66 | 15.00 | 30.58 | 40.90 | 82.37 | 88.83 | 75.74 | 0.21 | 9.92 |
| PIPPack (ensembled) | <u>0.386</u> | <u>0.289</u> | <u>0.475</u> | **8.76** | **13.94** | **28.69** | **38.47** | **84.05** | **90.27** | **77.60** | 0.14 | 13.91 |
| PIPPack+RS (ensembled) | 0.394 | 0.295 | 0.485 | <u>9.15</u> | <u>14.53</u> | <u>29.73</u> | 40.29 | <u>83.07</u> | <u>89.34</u> | 76.59 | 0.19 | <u>9.82</u> |

<sup>†</sup> The native Top2018 test set has 0.69% rotamer outliers and a clashscore of 1.27.
<sup>‡</sup> The mean values of three randomly seeded models are shown.

While PIPPack and DiffPack were trained to match the distribution of native χ angles, DLPacker and AttnPacker were trained to capture the distribution of atomic coordinates, and AttnPacker was trained specifically to minimize the RMSD between the predicted and native coordinates. DiffPack autoregressively captures conditional χ distributions via denoising one angle at a time, achieving strikingly better RMSD and clashscore even without post-processing. The confidence-aware DiffPack predicts multiple conformations and then selects regions with highest confidence (i.e., lowest predicted RMSD). Both DLPacker and Rosetta Packer consider residues one at a time, while both AttnPacker and PIPPack produce entire rotamers for each residue all at once. As mentioned by the authors^24^, the autoregressive nature of DiffPack and its ability for iterative refinement may contribute to the reduced clashes in output models and overall performance.

We next looked at the performance of each of these methods on a per amino acid basis. As shown in Figure 3 and Table S1, ensembled PIPPack improves the χ angle prediction in terms of rotamer recovery for most amino acid types (ARG, ASN, ASP, CYS, GLN, HIS, LEU, MET, PHE, SER, THR, TRP, and TYR) over the other PSCP methods, even DiffPack with confidence. Moreover, on individual χ_*i*_ basis, PIPPack demonstrates robust and competitive performance, achieving top-1 for 74% of all χ (Table S1). Ensembled PIPPack even produces improved RMSD for five amino acids (ASN, HIS, MET, SER, and TRP).

**Figure 3:**
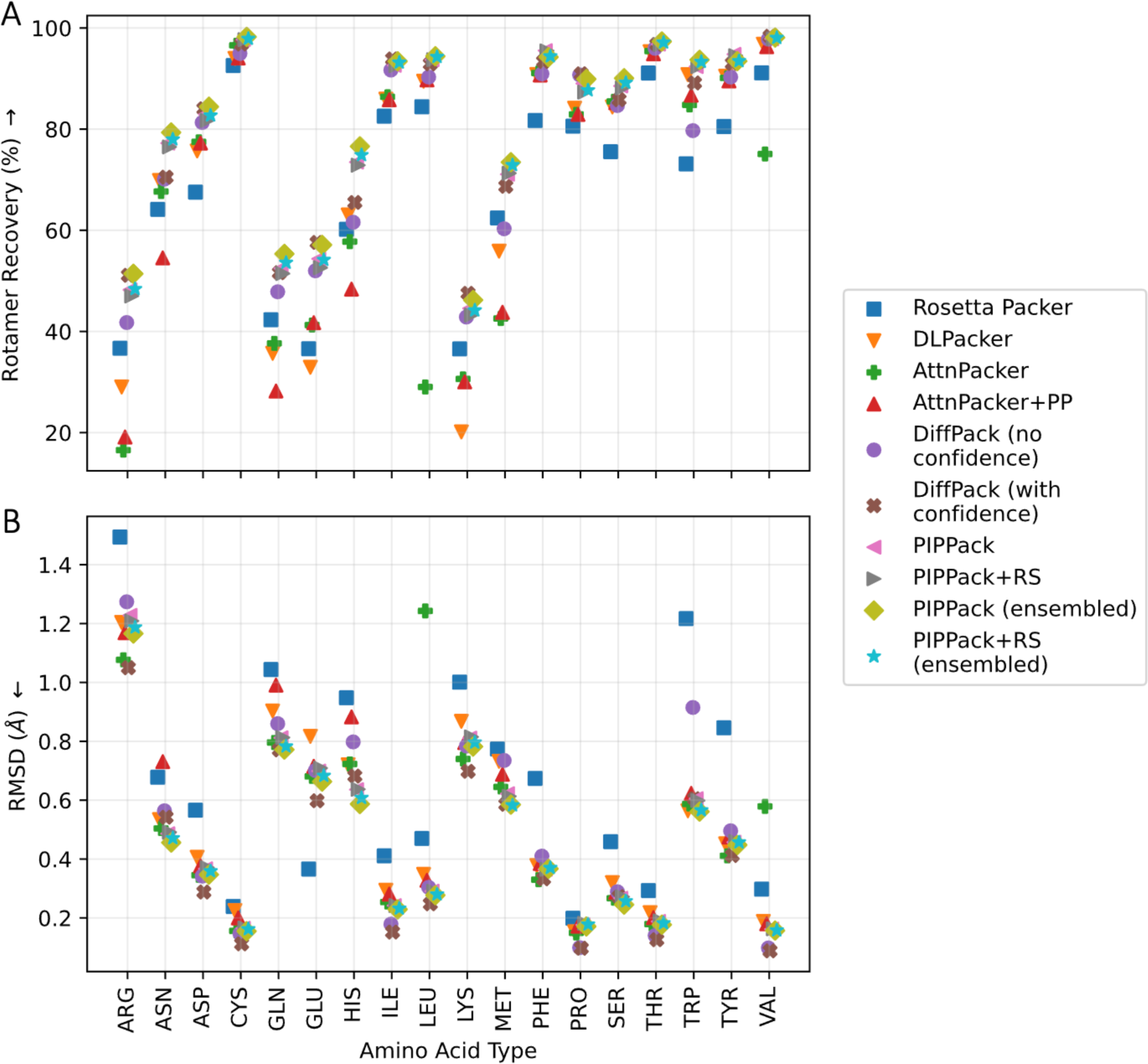
Side chain packing performance across amino acid type. (A) Ensembled PIPPack performs competitively with other PSCP methods in rotamer recovery, achieving top-1 performance for over half of all amino acid types (ARG, ASN, ASP, CYS, GLN, HIS, LEU, MET, PHE, SER, THR, TRP, and TYR). (B) With respect to RMSD, ensembled PIPPack also performs competitively for all amino acid types, even improving for certain residues (ASN, HIS, MET, SER, and TRP).

As PSCP is a crucial step in many computational workflows such as protein design and protein-protein docking, having rapid access to accurate side chains can dramatically impact the scale and performance of the simulations. To this end, we evaluated the runtimes of the various PSCP methods. While also being highly accurate, PIPPack additionally achieves the fastest runtimes (with and without post-prediction minimization) amongst the methods evaluated and for almost every protein size tested (Table 4). Moreover, because of the lightweight model and low resource demand, PIPPack can be run efficiently on both CPU and GPU.

**Table 4:** Runtime comparison between side chain packing methods.

| Method | Runtime<sup>†</sup> (sec / protein) |  |  |  |  |
| --- | --- | --- | --- | --- | --- |
| | $N_{res} = 100$ | $N_{res} = 250$ | $N_{res} = 500$ | $N_{res} = 750$ | $N_{res} = 1005$ |
| Rosetta Packer<sup>‡</sup> | 72.37 | 542.19 | 911.93 | 1836.12 | 2563.08 |
| DLPacker | 19.70 | 48.99 | 110.31 | 172.09 | 242.73 |
| AttnPacker | 1.70 | 2.24 | 3.91 | 6.20 | 8.82 |
| AttnPacker+PP | 7.78 | 12.65 | 14.21 | 16.74 | 21.10 |
| DiffPack | 5.98 | 7.42 | 11.27 | 15.66 | 21.36 |
| DiffPack +Confidence | 8.12 | 13.64 | 24.89 | 35.89 | 45.38 |
| PIPPack | 0.21 | 0.21 | 0.21 | 0.22 | 0.23 |
| PIPPack+RS | 0.64 | 0.63 | 0.65 | 0.73 | 0.73 |
| PIPPack (ensembled) | 0.65 | 0.63 | 0.63 | 0.64 | 0.69 |
| PIPPack+RS (ensembled) | 1.06 | 1.03 | 1.08 | 1.18 | 1.17 |

<sup>†</sup> Mean values of 25 unbatched predictions.
<sup>‡</sup> Only method solely running on the CPU.

### PIPPack Captures Complex Physical Interactions

Protein amino acid side chains are known to make many different types of interactions between one another and other biomolecules. These include the electrostatic interactions, van der Waals interactions, and hydrogen bonding, π-π stacking, and π-cation interactions, plus other less understood interactions. A desirable property of any PSCP method is the recapitulation of these interactions in the solutions that it produces. Physics-based methods, like Rosetta Packer, explicitly incorporates some of these interactions in specific score terms, such as van der Waals attractive and repulsive energies, hydrogen-bonding energy, and electrostatic energy. These interactions should also be able to be learned directly from the data, which is the assumption made by most DL-based methods including PIPPack. To investigate how well PIPPack can capture these complex physical interactions, we sought to find examples of several of these interactions. As shown in Figure 4, PIPPack reproduces van der Waals forces between well-packed hydrophobic residues (Fig. 4A), coordination of an “invisible ligand” by four cysteines (Fig. 4B), formation of a salt bridge between a lysine and an aspartic acid (Fig. 4C), hydrogen bonding between a serine and aspartate (Fig. 4D), π-π stacking interactions between aromatic rings (Fig. 4E), and π-cation interactions between an aromatic ring and a positively-charged lysine (Fig. 4F).

**Figure 4:**
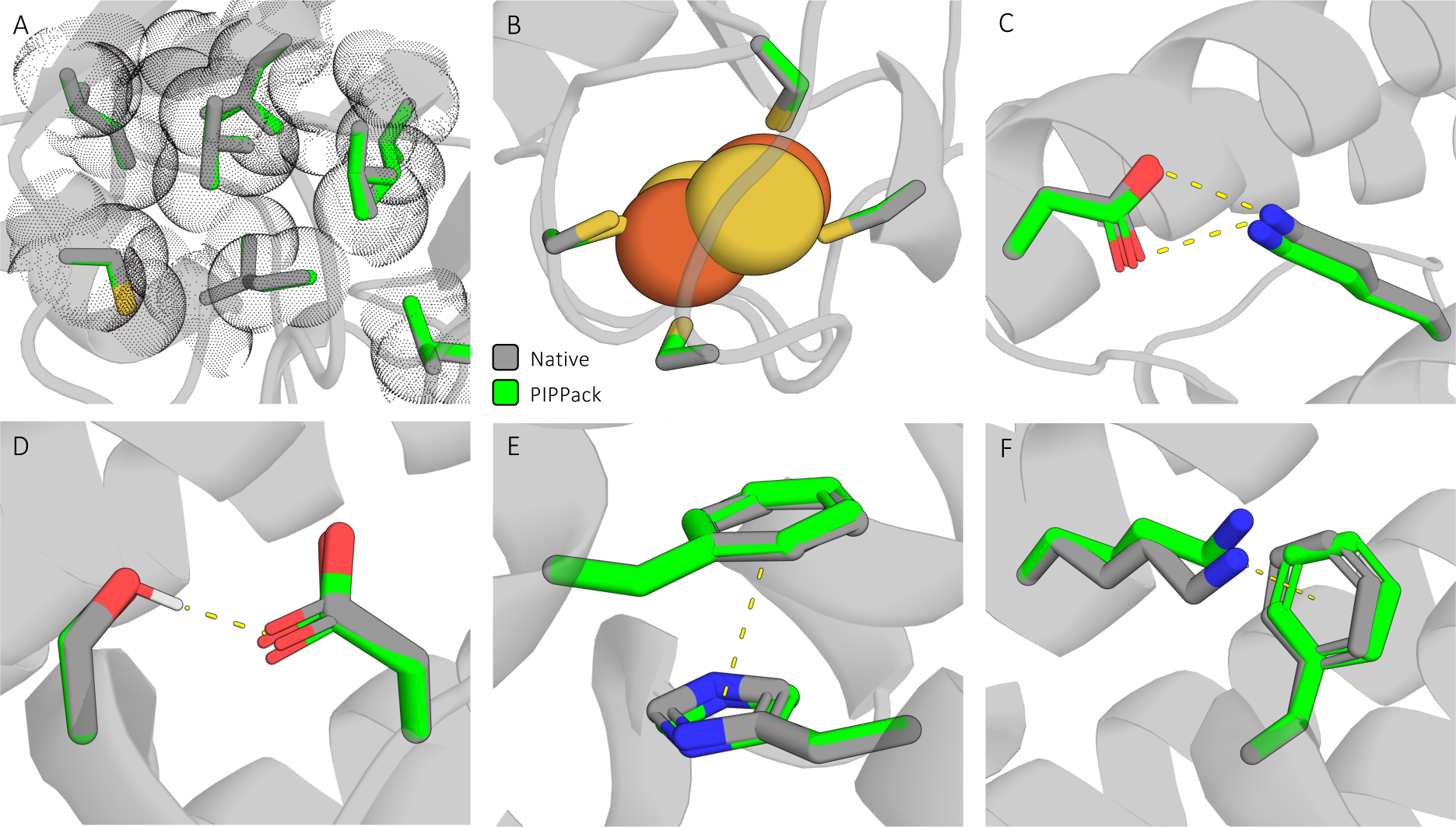
PIPPack reproduces physical interactions involving side chains. Numerous types of complex interactions are made by side chains within proteins, and PIPPack can reproduce many of them, including: (A) van der Waals forces, (B) coordination of invisible ligands, (C) ionic salt bridges, (D) hydrogen bonding, (E) π-π stacking, and (F) π-cation interactions.

## Discussion

Protein side chain packing is an important step in many computational protein simulations and can provide key insights into interactions and functional mechanisms. Because of its role in simulation, PSCP can heavily impact the performance of algorithms and, ideally, can provide rapid and accurate access to side chain conformations. Here, we present PIPPack which is an invariant GNN with novel message passing layers that has been trained to capture the native distribution of χ dihedral angles from native protein structures. Our model is the fastest among other state-of-the-art PSCP methods and produces competitive residue-level RMSDs and rotamer recovery, demonstrating its ability to recapitulate native side chain conformations.

Contributing to PIPPack’s success, we reframed the PSCP task as classification instead of regression, introduced iterative refinement via recycling, and developed a geometry-aware message passing scheme. The latter two were inspired by the success of the protein structure prediction network AF2. The novel message passing scheme, called invariant point message passing (IPMP), can be viewed as a generalization of AF2’s invariant point attention, as it can accommodate arbitrary residue neighborhoods and invariant message functions. Since the predictions from PIPPack are a probability distribution over χ angle bins, it is also possible to sample these distributions to generate ensembles of side chain conformations. Finetuning the model with auxiliary losses that act on a sample from the predicted distribution provides marginal benefits. The performance of our method is further bolstered by leveraging knowledge from multiple independently trained models in an ensemble.

PIPPack effectively captures the local environment of residues within a protein by propagating information along the protein graph, like AttnPacker^27^ and DiffPack^24^. In contrast, DLPacker^26^ voxelates the environment of each residue, performs convolutions to extract information, and sequentially assigns rotamers. Other DL-based PSCP methods have been announced, but we were unable to benchmark them with PIPPack because of unreleased code and/or model weights. OPUS-Rota4^23^ is a series of neural networks that processes local environmental features, evolutionary information in the form of a multiple sequence alignment, and the 3D-voxelized representation of the environment produced by DLPacker. ZymePackNet^28^ is a series of GNNs that build up the side chains of each amino acid by iteratively predicting χ angles given the partial context of the angles in the chain and then refines the previous predictions given the full context of the side chain.

We believe that since PIPPack produces rapid, accurate rotamer predictions, it will be a valuable resource that can speed up computational simulations without compromising on quality. Moreover, the generality of IPMP for protein representations may provide additional benefits for other protein-related tasks. Although PIPPack quickly generates reasonable predictions, it can still violate physical constraints through steric clashes. It remains future work to be able to reduce these clashes without secondary post-prediction optimization while also maintaining high accuracy.

## Conclusion

Protein side chains are responsible for the broad functions of proteins through their flexible interactions with each other and other biomolecules, highlighting the need for rapid and accurate protein side chain packing (PSCP) methods in *in silico* simulations and design. Here we present the Protein Invariant Point Packer (PIPPack) which utilizes a novel message passing scheme to learn high-quality distributions of the χ dihedral angles and outperforms other physics- and deep learning-based PSCP methods in rotamer recovery. Although reconstructing rotamers assuming ideality and iteratively refining its predictions through recycling, PIPPack still benefits from post-prediction optimization to reduce minor clashes, revealing a direction for future studies. Moreover, PIPPack does not consider any non-protein atoms when making its predictions, despite the obvious importance of modeling these interactions, suggesting another route for improvement.

## Acknowledgements

This work would not have been possible without the enduring love and support from Scar and their cats Oracle and Sushi. We also would like to thank the rest of the Kuhlman lab for their insightful discussions throughout this project. This work was supported by the NIH grant R35GM131923 (B.K.) and by the NSF fellowship DGE-2040435 (N.Z.R.).

## Data availability

The training and inference code for the method developed in this study are openly available in the PIPPack GitHub repository at https://github.com/Kuhlman-Lab/PIPPack.

---

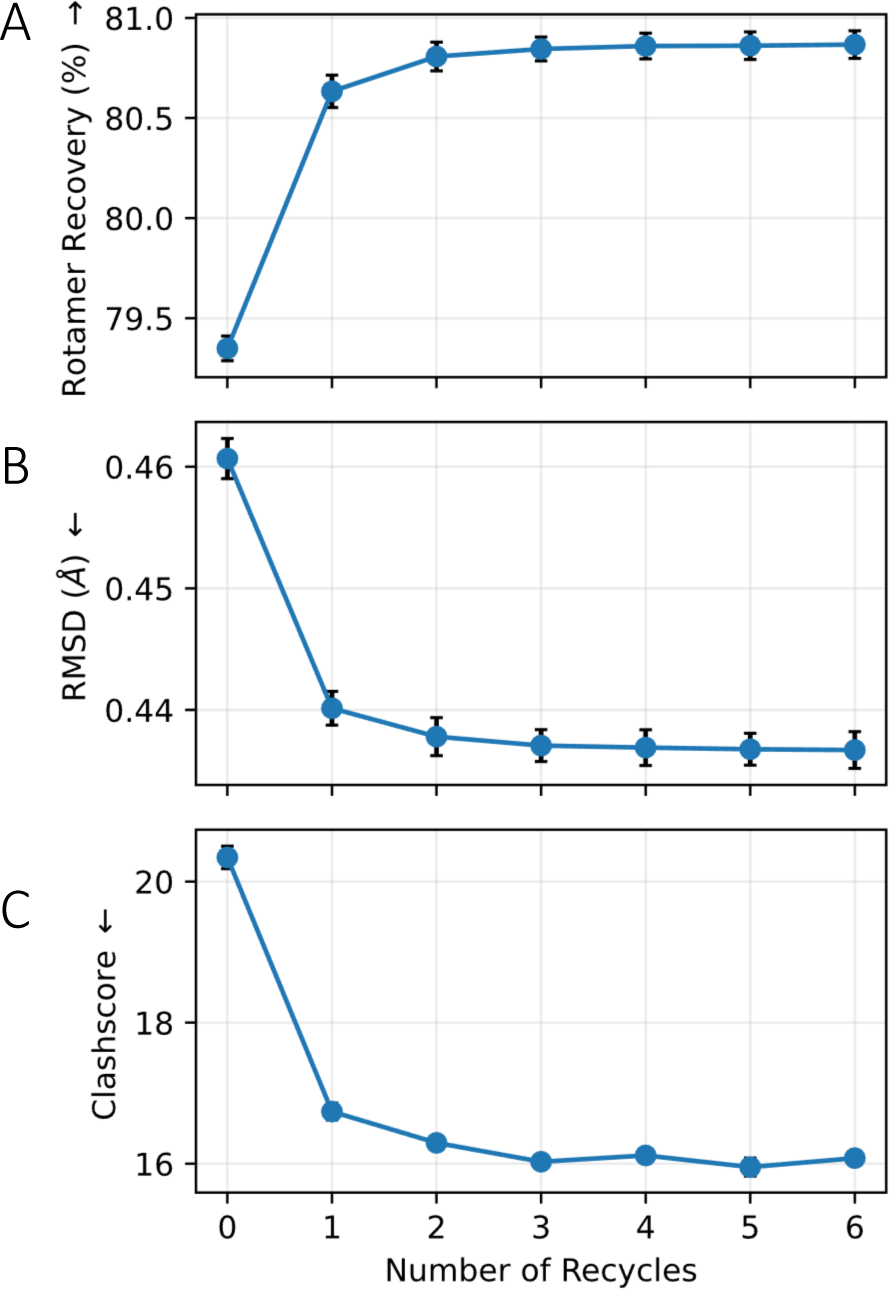

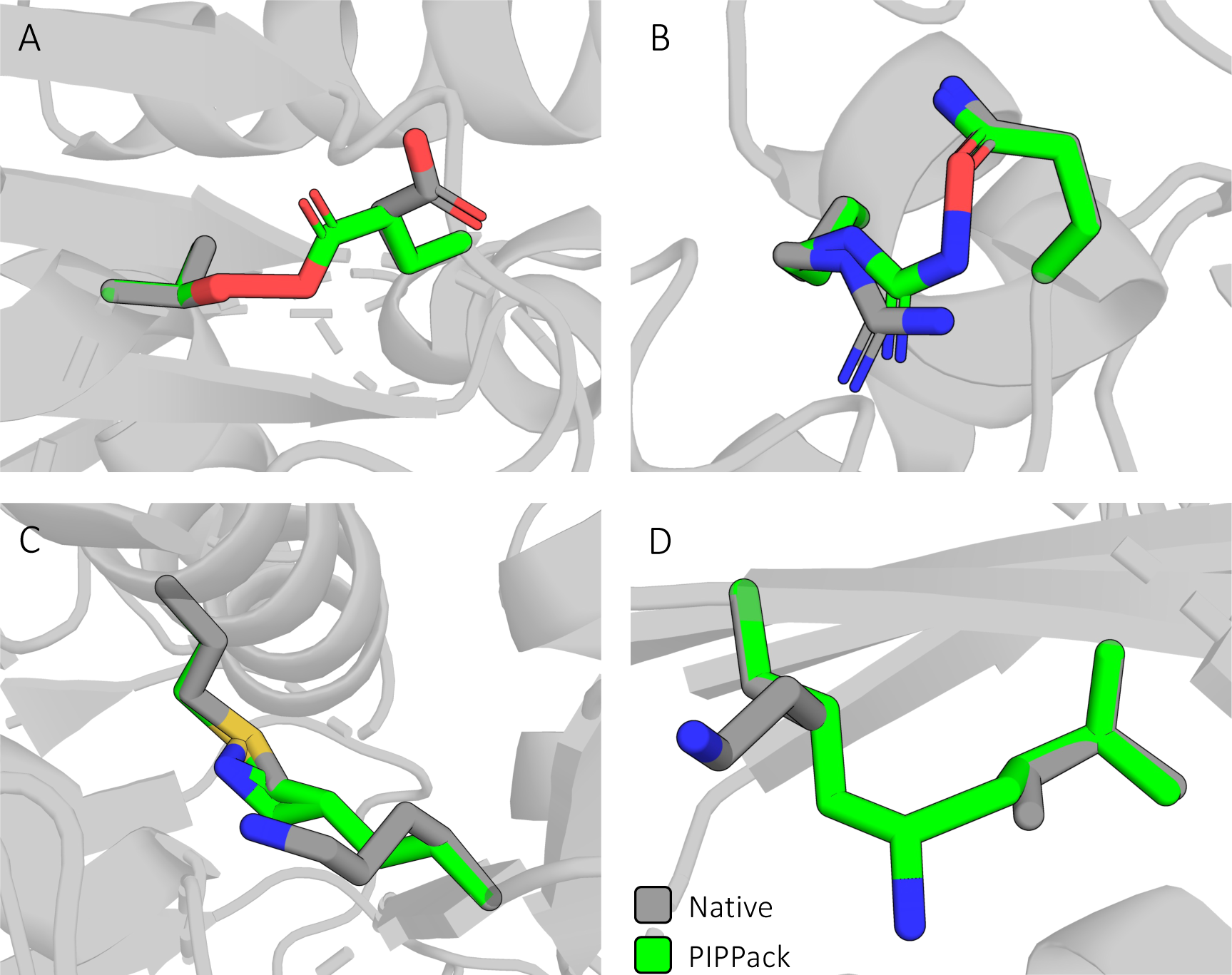

