## Supplemental Materials for "Invariant point message passing for protein side chain packing"

**TITLE:**

**RUNNING TITLE:**

Protein side chain packing

Nicholas Z. Randolph[1,2] and Brian Kuhlman[1,2]

[1]Department of Bioinformatics and Computational Biology, University of North Carolina School of Medicine, Chapel Hill, North Carolina, USA.

[2]Department of Biochemistry and Biophysics, University of North Carolina School of Medicine, Chapel Hill, North Carolina, USA.

**Table of Contents** **Page**

| | |
|---|---|
| Figure S1: PIPPack performance across recycles . | 3 |
| Figure S2: Representative Clashes Produced by PIPPack. | 4 |
| Table S1: Side chain $\chi$ angle errors across amino acid type. | 5 |
| Table S2: Rotamer Evaluations for PSCP Methods. | 10 |

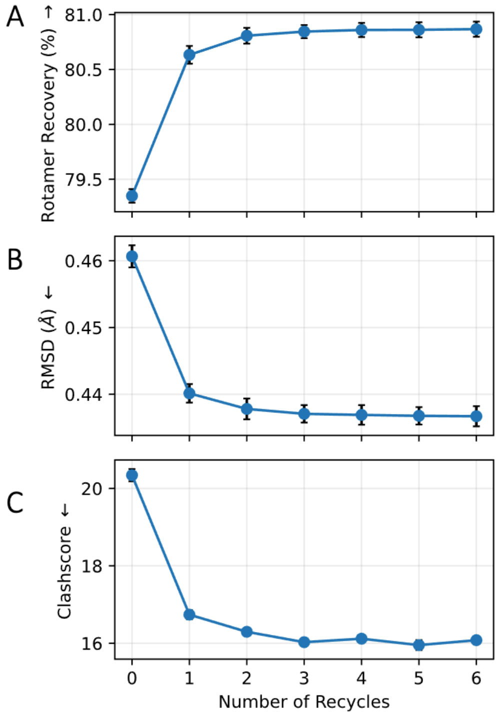

**Figure S1: PIPPack performance across recycles.** PIPPack uses recycling to iteratively refine its side chain conformation predictions, resulting in improved performance metrics. Recycling past the value PIPPack was trained for (in our case 3) does not result in any significant differences in performance, but the model performs well even with a single recycle.

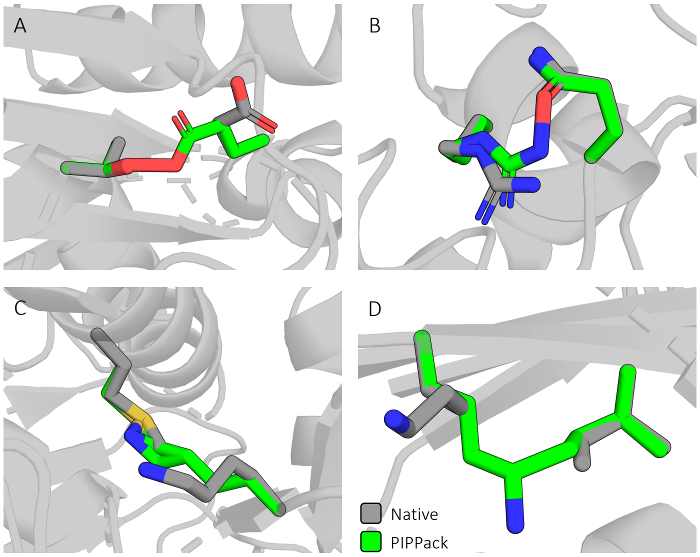

**Figure S2: Representative Clashes Produced by PIPPack**. Although not common, PIPPack can produce atomic clashes due to the one-shot nature of the network. Egregious clashes, as shown here, are usually fixed via post-prediction minimization. Most of these clashes occur between hydrogen bonding partners (A and B) and/or long side chains (B, C, and D).

### Table S1: Side chain $\chi$ angle errors across amino acid type.

| AA Type | Method | MAE (°) ↓ | | | | AA Type | Method | MAE (°) ↓ | |
|---|---|---|---|---|---|---|---|---|---|
| | | $\chi_1$ | $\chi_2$ | $\chi_3$ | $\chi_4$ | | | $\chi_1$ | $\chi_2$ |
| **ARG** | Rosetta Packer | 23.00 | 24.34 | 48.62 | 48.54 | **ASN** | Rosetta Packer | 20.35 | 28.72 |
| | DLPacker | 17.11 | 22.76 | 45.97 | 53.64 | | DLPacker | 13.19 | 28.56 |
| | AttnPacker | 15.61 | 20.11 | 43.56 | 48.96 | | AttnPacker | 12.07 | 27.69 |
| | AttnPacker+PP | 17.22 | 21.08 | 42.78 | 48.56 | | AttnPacker+PP | 12.36 | 53.56 |
| | DiffPack | 16.50 | 19.86 | 41.19 | 46.56 | | DiffPack | 14.29 | 33.27 |
| | DiffPack +Confidence | 13.30 | 17.18 | 34.86 | **40.30** | | DiffPack +Confidence | 11.74 | 36.16 |
| | PIPPack† | 13.60 | 17.79 | 36.15 | 42.36 | | PIPPack† | 11.24 | 23.66 |
| | PIPPack+RS† | 13.90 | 18.03 | 36.67 | 42.34 | | PIPPack+RS† | 11.24 | 24.42 |
| | PIPPack (ensembled) | **12.62** | **16.46** | **33.98** | 39.79 | | PIPPack (ensembled) | **10.22** | **22.06** |
| | PIPPack+RS (ensembled) | 13.54 | 17.61 | 35.97 | 41.68 | | PIPPack+RS (ensembled) | 10.66 | 23.17 |
| **LYS** | Rosetta Packer | 23.40 | 25.05 | 32.14 | 41.89 | **ASP** | Rosetta Packer | 19.72 | 18.1 |
| | DLPacker | 17.03 | 24.37 | 38.45 | 60.48 | | DLPacker | 12.86 | 15.40 |
| | AttnPacker | 14.77 | 20.50 | 30.16 | 41.27 | | AttnPacker | 10.40 | 14.09 |
| | AttnPacker+PP | 15.64 | 21.08 | 30.50 | 41.37 | | AttnPacker+PP | 10.80 | 13.96 |
| | DiffPack | 15.73 | 19.67 | 28.48 | 40.06 | | DiffPack | 11.95 | 12.67 |
| | DiffPack +Confidence | 13.35 | **17.83** | **25.73** | 38.45 | | DiffPack +Confidence | **9.59** | 11.55 |
| | PIPPack† | 14.09 | 19.22 | 26.46 | 38.09 | | PIPPack† | 10.44 | 11.92 |
| | PIPPack+RS† | 14.20 | 19.50 | 26.85 | 39.28 | | PIPPack+RS† | 10.55 | 12.34 |
| | PIPPack (ensembled) | **13.12** | 18.23 | 25.76 | **36.99** | | PIPPack (ensembled) | **9.59** | **11.28** |
| | PIPPack+RS (ensembled) | 13.59 | 18.84 | 26.41 | 38.71 | | PIPPack+RS (ensembled) | 10.03 | 11.95 |
| **GLU** | Rosetta Packer | 29.33 | 33.46 | 27.00 | | **LEU** | Rosetta Packer | 12.64 | 18.53 |

| | | | | | | | | | |
|---|---|---|---|---|---|---|---|---|---|
| | DLPacker | 23.15 | 29.94 | 30.35 | | | DLPacker | 8.33 | 14.25 |
| | AttnPacker | 18.82 | 24.44 | 26.82 | | | AttnPacker | 7.13 | 87.63 |
| | AttnPacker+PP | 19.34 | 24.57 | 26.36 | | | AttnPacker+PP | 7.36 | 13.44 |
| | DiffPack | 19.55 | 25.16 | 22.54 | | | DiffPack | 9.57 | 11.77 |
| | DiffPack +Confidence | 16.35 | 21.31 | 20.48 | | | DiffPack +Confidence | 7.33 | 9.49 |
| | PIPPack† | 17.48 | 22.94 | 21.49 | | | PIPPack† | 6.18 | 9.71 |
| | PIPPack+RS† | 17.79 | 23.33 | 22.06 | | | PIPPack+RS† | 6.19 | 9.76 |
| | PIPPack (ensembled) | **16.33** | **21.24** | **20.41** | | | PIPPack (ensembled) | **5.70** | **9.00** |
| | PIPPack+RS (ensembled) | 16.86 | 22.27 | 21.41 | | | PIPPack+RS (ensembled) | 5.77 | 9.18 |
| **GLN** | Rosetta Packer | 23.20 | 32.63 | 42.28 | | **ILE** | Rosetta Packer | 12.48 | 19.03 |
| | DLPacker | 18.17 | 27.87 | 43.74 | | | DLPacker | 6.86 | 17.78 |
| | AttnPacker | 15.67 | 24.47 | 42.01 | | | AttnPacker | 6.06 | 14.92 |
| | AttnPacker+PP | 16.29 | 24.68 | 71.75 | | | AttnPacker+PP | 6.43 | 15.07 |
| | DiffPack | 16.95 | 24.76 | 46.42 | | | DiffPack | 4.66 | 11.52 |
| | DiffPack +Confidence | 13.99 | 21.43 | 46.27 | | | DiffPack +Confidence | **4.02** | **9.64** |
| | PIPPack† | 14.25 | 22.07 | 39.36 | | | PIPPack† | 5.01 | 10.93 |
| | PIPPack+RS† | 14.24 | 22.13 | 40.14 | | | PIPPack+RS† | 5.03 | 11.02 |
| | PIPPack (ensembled) | **13.23** | **20.35** | **37.88** | | | PIPPack (ensembled) | 4.63 | 10.07 |
| | PIPPack+RS (ensembled) | 13.54 | 21.16 | 38.80 | | | PIPPack+RS (ensembled) | 4.71 | 10.34 |
| **MET** | Rosetta Packer | 17.77 | 21.39 | 39.55 | | **PRO** | Rosetta Packer | 10.90 | 15.14 |
| | DLPacker | 13.61 | 20.02 | 41.74 | | | DLPacker | 8.45 | 12.42 |
| | AttnPacker | 12.16 | 17.52 | 43.65 | | | AttnPacker | 8.10 | 11.74 |
| | AttnPacker+PP | 13.02 | 17.90 | 43.06 | | | AttnPacker+PP | 8.43 | 12.28 |
| | DiffPack | 15.53 | 20.05 | 40.67 | | | DiffPack | 5.28 | 6.35 |

| | | | | | | | | | |
|---|---|---|---|---|---|---|---|---|---|
| | DiffPack +Confidence | 11.55 | 15.30 | 32.51 | | | DiffPack +Confidence | **5.17** | **6.20** |
| | PIPPack† | 10.25 | 13.40 | 30.90 | | | PIPPack† | 6.97 | 9.24 |
| | PIPPack+RS† | 10.33 | 13.59 | 30.44 | | | PIPPack+RS† | 7.52 | 10.18 |
| | PIPPack (ensembled) | **9.46** | **11.95** | **28.66** | | | PIPPack (ensembled) | 6.56 | 8.62 |
| | PIPPack+RS (ensembled) | 9.69 | 12.72 | 28.76 | | | PIPPack+RS (ensembled) | 7.28 | 9.89 |
| | Rosetta Packer | 13.14 | 11.70 | | | | Rosetta Packer | 14.46 | |
| | DLPacker | 6.45 | 8.85 | | | | DLPacker | 10.05 | |
| | AttnPacker | 5.26 | 8.56 | | | | AttnPacker | 8.15 | |
| | AttnPacker+PP | 5.81 | 8.59 | | | | AttnPacker+PP | 8.44 | |
| | DiffPack | 8.32 | 7.56 | | | | DiffPack | 7.78 | |
| **PHE** | DiffPack +Confidence | 6.49 | 6.50 | | | **THR** | DiffPack +Confidence | 6.91 | |
| | PIPPack† | 5.12 | **5.99** | | | | PIPPack† | 7.31 | |
| | PIPPack+RS† | 5.12 | 6.07 | | | | PIPPack+RS† | 7.50 | |
| | PIPPack (ensembled) | **4.71** | 8.00 | | | | PIPPack (ensembled) | **6.61** | |
| | PIPPack+RS (ensembled) | 4.84 | 7.75 | | | | PIPPack+RS (ensembled) | 6.93 | |
| | Rosetta Packer | 14.98 | 11.63 | | | | Rosetta Packer | 32.99 | |
| | DLPacker | 7.06 | 8.82 | | | | DLPacker | 22.08 | |
| | AttnPacker | 5.99 | 8.92 | | | | AttnPacker | 18.20 | |
| | AttnPacker+PP | 6.39 | 8.96 | | | | AttnPacker+PP | 18.46 | |
| **TYR** | DiffPack | 8.99 | 7.47 | | | **SER** | DiffPack | 21.53 | |
| | DiffPack +Confidence | 7.32 | 6.62 | | | | DiffPack +Confidence | 20.17 | |
| | PIPPack† | 5.81 | **6.23** | | | | PIPPack† | 17.03 | |
| | PIPPack+RS† | 5.92 | 6.45 | | | | PIPPack+RS† | 17.51 | |

| | | | | | | | | |
|---|---|---|---|---|---|---|---|---|
| | PIPPack (ensembled) | **5.29** | 8.13 | | | | PIPPack (ensembled) | **15.19** |
| | PIPPack+RS (ensembled) | 5.60 | 7.71 | | | | PIPPack+RS (ensembled) | 16.33 |
| **TRP** | Rosetta Packer | 17.63 | 30.62 | | | **CYS** | Rosetta Packer | 11.73 |
| | DLPacker | 6.99 | 14.97 | | | | DLPacker | 10.48 |
| | AttnPacker | 6.84 | 15.76 | | | | AttnPacker | 6.83 |
| | AttnPacker+PP | 7.11 | 15.39 | | | | AttnPacker+PP | 8.58 |
| | DiffPack | 10.53 | 26.73 | | | | DiffPack | 8.27 |
| | DiffPack +Confidence | 8.30 | 14.77 | | | | DiffPack +Confidence | 6.14 |
| | PIPPack† | 6.25 | 12.13 | | | | PIPPack† | 5.94 |
| | PIPPack+RS† | 6.33 | 12.07 | | | | PIPPack+RS† | 6.02 |
| | PIPPack (ensembled) | **5.51** | **10.57** | | | | PIPPack (ensembled) | **5.23** |
| | PIPPack+RS (ensembled) | 5.95 | 10.79 | | | | PIPPack+RS (ensembled) | 5.75 |
| **HIS** | Rosetta Packer | 18.61 | 45.62 | | | **VAL** | Rosetta Packer | 14.02 |
| | DLPacker | 10.48 | 47.03 | | | | DLPacker | 7.54 |
| | AttnPacker | 9.54 | 52.95 | | | | AttnPacker | 32.93 |
| | AttnPacker+PP | 10.32 | 69.95 | | | | AttnPacker+PP | 7.18 |
| | DiffPack | 12.77 | 53.06 | | | | DiffPack | 5.21 |
| | DiffPack +Confidence | 9.74 | 49.34 | | | | DiffPack +Confidence | **4.62** |
| | PIPPack† | 8.95 | 37.93 | | | | PIPPack† | 5.83 |
| | PIPPack+RS† | 8.98 | 38.17 | | | | PIPPack+RS† | 5.85 |
| | PIPPack (ensembled) | **8.05** | **34.37** | | | | PIPPack (ensembled) | 5.47 |

| | | | | | | | | |
|---|---|---|---|---|---|---|---|---|
| | PIPPack+RS (ensembled) | 8.51 | 35.90 | | | | PIPPack+RS (ensembled) | 5.56 |

**Table S2: Rotamer Evaluations for PSCP Methods.**

| Method | Rotamer Evaluation (%) | | |
|---|---|---|---|
| | Favored | Allowed | Outlier |
| Native | 96.77 | 2.54 | 0.69 |
| Rosetta Packer | 99.29 | 0.65 | 0.07 |
| DLPacker | 96.33 | 2.50 | 1.17 |
| AttnPacker | 87.65 | 8.34 | 4.01 |
| AttnPacker+PP | 93.62 | 4.09 | 2.29 |
| DiffPack | 97.97 | 1.48 | 0.55 |
| DiffPack +Confidence | 98.35 | 1.23 | 0.43 |
| PIPPack | 98.93 | 0.90 | 0.17 |
| PIPPack+RS | 98.71 | 1.08 | 0.21 |
| PIPPack (ensembled) | 99.03 | 0.82 | 0.14 |
| PIPPack+RS (ensembled) | 98.78 | 1.03 | 0.19 |
